# Assessing the Relative Impact of Grasp and Object on Inferior Frontal Gyrus Activity during a Grasping Task

**DOI:** 10.64898/2026.06.30.735663

**Authors:** Emily C. Conlan, Crispin Foli, William D. Memberg, Eric Z. Herring, Jennifer A. Sweet, A. Bolu Ajiboye

## Abstract

The lateral grasp network, responsible for translating visual properties of an object to execution of a motor act, is comprised of the anterior intraparietal area (AIP), area F5, and the primary motor cortex (M1). Non-human primate studies of F5 have shown that it encodes a wide range of hand positions and object properties. Human studies in F5’s human homologue, the inferior frontal gyrus (IFG), have leveraged this area’s ability to encode grasp-object pairs for the purposes of Brain Machine Interface (BMI) control. However, whether modulation is driven by grasp, object, or the interaction between grasp and object is unclear. In the present study, intracortical IFG signals were recorded from human participants with tetraplegia during a motor visualization task where grasp and object were varied. Grasp was found to be the predominant factor driving modulation of IFG signals. Object was found to only be weakly represented in neural data. However, object contribution peaked earlier than grasp contribution, indicating early integration of object information. Grasp-object interactions were also found to have a significant impact. Cortical separation between object conditions varied based on the grasp used. In addition, subspace analysis showed that the underlying neural population structure associated with each grasp type was significantly different from one another. Despite the impact of grasp-object interactions, the present study demonstrates that grasp is the primary driver of inferior frontal gyrus neural modulation.

## Introduction

The lateral grasp network [1] identifies three key cortical areas implicated in the transformation of an object’s properties into grasp execution. The anterior intraparietal area (AIP) encodes visual features of objects that can be used to guide grasping (called object affordances). AIP is densely connected to area F5, which receives information about object affordances and translates them into potential motor acts [2], or possible grasping actions that could be used to interact with the object features derived from AIP. In addition, F5 is multipurpose and encodes both object and motor properties during movement preparation [3]. Finally, F5’s connection with primary motor cortex (M1) translates the potential motor act into execution of movement.

The multifunctional role of F5 is due to the presence of both motor and visuomotor neurons. Motor neurons are only active during motor execution regardless of the object presented [4], whereas visuomotor neurons (mirror and canonical [5], [6]), are generally active during visual observation of a motor act. Mirror neurons activate similarly when executing a hand grasp and when observing someone else perform the action. These neurons are thought to aid in translating an observed act into motor function commands that could be used for the purposes of imitation, action understanding, and communication [7]. Canonical neurons in F5 discharge during grasping action or observation of an object. These neurons form a library or motor dictionary that is responsive to objects or grasp movements fulfilling specific parameters, such as object size, shape, or orientation. These visuomotor type neurons discharge during object observation to encode the potential motor act, regardless of whether the grasp will be made [4], [8]. Visuomotor neurons have been used to classify grasp type before movement occurs with greater than 95% classification accuracy [9].

Transcranial magnetic stimulation (TMS) and functional magnetic resonance imaging (fMRI) studies have shown that F5’s human homologue, IFG, shares similar properties. From fMRI studies, ventral premotor cortex, parietal, motor, and sensory regions are active during object manipulation [10]. Virtual lesions induced using TMS resulted in impaired grasping by either disrupting the proper recruitment of intrinsic hand muscles or finger positioning on an object [11]. Few groups have investigated the function of IFG in human living with tetraplegia to characterize how it encodes different grasp types. One study investigated IFG activity during presentation of specific grasp-object pairings [12], showing condition-dependent single unit modulation during the cue (premovement) phase of grasp. However, it is unclear in this study if the observed cortical activity was driven only by the grasp type presented or due to the coupling of grasp and object. In the present study, we explored how IFG responds to grasp and object independently, as well as the interaction between grasp and object. Since the primary function of this area in NHP and human studies is to create an appropriate grasp shape for the object presented, we hypothesized that grasp will be the largest factor driving modulation of IFG neural signals. Further, due to the presence of visuomotor (specifically canonical) neurons in IFG, we hypothesized that the contribution of object and the interaction between grasp and object would be non-negligible to IFG neural modulation.

## Methods

### Participant Details

A 27-year-old right-handed male (ID number RP1) with C3-C4 level American Spinal Injury Association Impairment Scale B (AIS-B) tetraplegia resulting in motor complete and sensory incomplete loss was enrolled into the Reconnecting Hand and Arm to Brain (ReHAB) Clinical Trial (ClinicalTrials.gov ID: NCT03898804). This research was conducted under an Investigational Device Exemption from the U.S. Food and Drug Administration and approved by the local Institutional Review Board. Informed consent was obtained from the participant prior to enrollment. The participant was implanted with six 8 x 8 (64-channel) 1.5-mm iridium oxide microelectrode arrays (Blackrock Microsystems, Salt Lake City, UT) in the left hemisphere in January 2021. Two arrays were implanted in primary motor cortex (M1), two in primary sensory cortex (S1), one in the anterior intraparietal area (AIP), and one in the inferior frontal gyrus (IFG) as shown in **Figure 1a**. Further details of the implanted system were previously reported [13].

**Figure 1:**
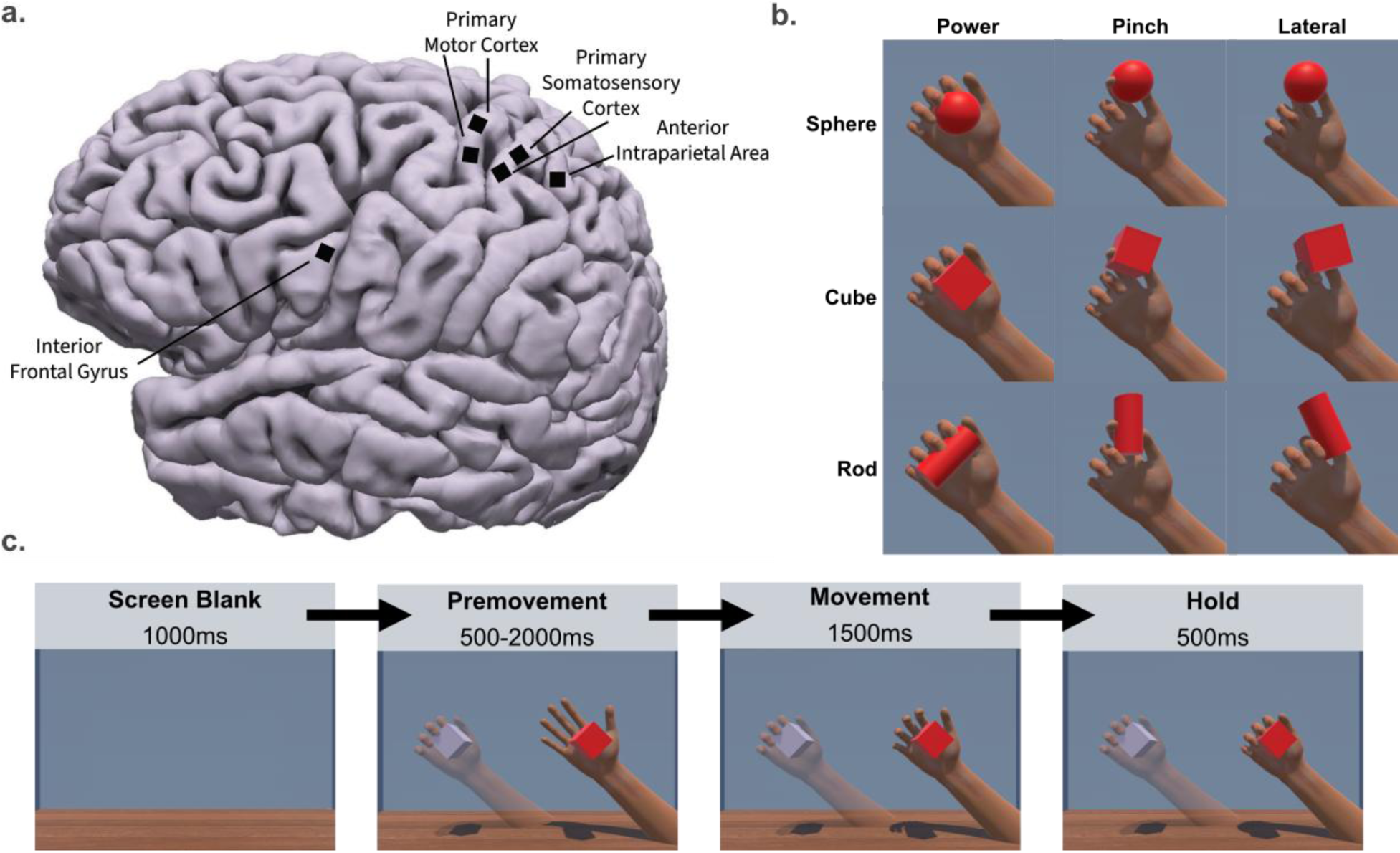
Implantation Details and Task Overview. a) Participant RP1 was implanted with six 8x8 microelectrode arrays. Two were implanted in primary motor cortex, two in primary somatosensory cortex, one in the anterior intraparietal area, and one in the inferior frontal gyrus. b) Three objects and three grasps were presented in the motor imagery task resulting in nine experimental conditions. A randomized grasp-object pair was selected for each trial. c) Each trial consisted of four epochs: screen blanking, premovement, movement, and hold. Due to the variability in the premovement epoch, only the first 1500ms were used in analysis.

### Virtual Task

A motor imagery visualization was presented to the participant in blocks of 5 minutes where 30-35 trials of hand grasping were performed. The participant was instructed to attempt to make the movements presented to him on screen. In the visualization, a random grasp-object pair was chosen from a set of nine possible combinations (three grasps and three objects) every trial. The grasps presented were power, pinch, and lateral or key grasp, and the object presented were a sphere, cube, or rod (**Figure 1b**). Every trial contained four epochs (**Figure 1c**): screen blank where nothing is displayed in the visualization space (1500ms), premovement (1500-2000ms), movement (1500ms), and hold (500ms). During the premovement epoch, the grasp-object pair was displayed to the left of the movable hand while the movable hand on the right remained in a flat resting position. During the movement phase, the right hand moved from the rest position to the intended grasping condition over the object. The position of the hand, size of the object, and the aperture of the grasp remained consistent across all nine grasp-object pairs.

The location and rotation of the object in virtual space changed so that the hand could remain in the same position on the screen while still executing the nine grasp-object combinations but was consistent within grasp types. Seven days of data were collected resulting in 35-50 trials per condition.

### Cortical Recordings

Data was recorded from all six arrays simultaneously at 30k samples/second. Data was band pass filtered (250-5,000 Hz), common average referenced and down sampled to 1ms samples to minimize offline data processing load. Spike band power (SBP, 250-5000Hz) was selected for decoding based on previous research demonstrating that SBP modulation encodes kinematic parameters of the hand [14]. Trial smoothing was performed by convolving block data with a gaussian curve (σ=50m). Noisy data was triaged by calculating the maximum value of each trial across all trials and all features. If a single trial’s maximum value was outside of the distribution of maximum values for other trials/features for more than 10% of features, the trial was eliminated. A window of activity was selected for each epoch for further analysis. 1000ms before cue onset to cue presentation was selected as the screen blanking epoch.

1500ms after cue presentation was selected for the premovement epoch, 1500ms after the movement cue was selected as the movement epoch, and 500ms after the end of movement was selected as the hold epoch. Each trial was then z-scored by the 1000ms screen blanking epoch across all trials from that specific day to generate normalized spike band power. Data was then concatenated across days for data analysis.

### Data Analysis

#### I. Single Feature Analysis

For each SBP feature, the area under the curve of the normalized spike band power traces was calculated in each epoch. All channels were categorized as: non-modulating, condition independent, or condition-dependent. An initial one-way analysis of variance (ANOVA) identified features that modulated from baseline, regardless of condition. Screen blanking epoch data was compared to each epoch of interest (premovement, movement) to identify modulating vs. non-modulating features. Only one of the nine conditions needed to be found significantly different from the screen blanking epoch data for SBP to be considered modulating. Modulating features were then categorized as condition-dependent or condition independent using a 2-way ANOVA with post-hoc multiple comparisons corrections using the Tukey-Kramer method. Features that were found to modulate to grasp, object, or the interaction between grasp and object and had a significant difference between at least one pair of conditions were classified as condition-dependent. All other features were classified as condition independent. Condition-dependent modulating features were further broken down into grasp-modulating, object-modulating, and/or interaction modulating features based on the results of the post-hoc Tukey-Kramer corrections.

The contribution of grasp, object, and the interaction between grasp and object were evaluated for all features regardless of modulation category. Using the omega-squared (ω²) metric each feature was evaluated for each factor’s overall impact on the feature data where: ω² < 0.01: “negligible contribution”, 0.01 < ω² < 0.06: “small contribution”, 0.06 < ω² < .14: “medium contribution”, ω² ≤ 0.14: “large contribution” [15]. Omega-squared is a measurement of the effect size of the factors in a dataset and is calculated as follows:

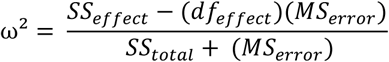

Where the SS is the Sum of Squares value as reported in the results of a 2-way ANOVA for the effect, or the factors in the dataset, *df_effect_* is the degrees of freedom within each factor, and *MS_error_* is the mean square variance of the error term. In essence, this variable is comparable to *R*^2^ for regression models. As data was concatenated across days, day’s individual contribution as well as the interaction between day and the data’s other factors (day: object, day: grasp, day : grasp : object) was calculated, but omitted from plots. The day contributions can be found in the **Supplementary Figure 1**.

#### II. Population Analysis

Prior to population level analyses, each day was aligned to the first day of data collection (reference day) to account for general neural drift due to variable neuron recording [16], [17], [18] .This was achieved by using a modified version of the alignment method outlined in Gallego et. al. [18]. In that method, an equal number of trial counts per day were extracted from all trials recorded so that each day had an equal number of trial counts. The two data sets were then projected into reduced dimensional space to reveal the latent dimensions across all trials. However, due to the variability in recording time, a largely variable number of trials per condition were recorded each day. Therefore, instead of finding the latent dimensions across all trials, data was trial averaged across the nine conditions prior to alignment resulting in a C x T x N matrix where C is the number of condition, T is the number of time points, and N is the number of features. This matrix was then unrolled into a C*T x N matrix. PCA was applied to the unrolled matrix to find the top ten latent factors in the condition averaged data for each recording day. Each day’s data was aligned to the reference data through a Procrustes analysis [17] which, unlike other methods such as CCA, allows the option to omit rotations and translations. Translations were omitted as block-to-block baseline drift was already removed via block-wise z-scoring, while rotations were omitted to preserve the fixed physical spatial identity of individual electrode channels and prevent artificial over-alignment of neural manifolds. The alignment between day X and the reference day was then applied to individual trial data resulting in the original number of trials, but with a reduction in features from the original 64 to those representing our latent dimension of 10. The validation of the alignment procedure can be found in **Supplementary Figure 2**. The following population level analysis techniques were used to identify underlying patterns of activity across all recorded channels [19].

#### A. Modulation Depth

Modulation depths [20] were calculated as the average Euclidean distance from baseline for each experiment condition using a jackknife approach. In addition, Euclidean distance was calculated between pairs of conditions to assess the overall population structure of the data. The smaller the distance, the closer conditions are to one another in 64 dimensional space, and hence the more similar their cortical representations. Euclidian distances above the diagonal were then categorized into within-grasp, outside-grasp, within-object, or outside-object Since each group could have a different number of data points, a permutation test was used to generate 1,000 iterations of a null sample to determine statistical significance between groups.

#### B. Linear Discriminant Analysis Classification

Linear discriminant analysis (LDA) was used to determine cross-validated decoding accuracy.

Classification accuracy was determined via a 5-fold cross-validated linear discriminant model. To account for the variable number of trials per condition, the LDA was run in 1,000 iterations with a random selection of trials each time. Cross validated decoding accuracy was determined as the average result over the 1000 iterations.

#### C. Time-Varying Principal Component Analysis

Principal component analysis (PCA) was performed to assess the temporal progression of the population response based on the methods described in Willet et. al. 2020 [20]. A matrix of data with dimensions N (number of neurons) x T (number of time steps) x R (number of trials) x P (pair types) was built and then trial averaged over pair type. The resulting N x T x P matrix type was unrolled into a N x TP matrix. In the original method proposed by Willet et. al. PCA was then applied to this matrix to determine the first ten principal components of the data using all trials. However, since the data has already been projected onto the latent factors, we omitted applying PCA in this step. A bootstrapping procedure was then used to build the N x TP matrix using a random selection of the same number of trials as the original data set with replacement. The 1000 iterations of bootstrapping was used to calculate confidence intervals and significant difference between individual pair trajectories.

#### D. Subspace Alignment

Principal component analysis was also performed to assess the overlap in low dimensional projections of grasping activity by object type. Here, PCA was used to obtain the top 5 principal components, or manifold, of sphere, cube, and rod data. To assess how well the neural activity of one object type is captured by another object type, the neural data of one object type was projected onto another object type’s manifold (i.e. projecting sphere data onto the cube manifold). Variance Accounted For (VAF) was calculated and compared to the VAF when projecting into the same object manifold. Similar to the above calculation of the time varying projections, the N x T x R x P matrix was averaged over a random sampling of trials within each pair type. The resulting N x T x P matrix type was unrolled into a N x TP matrix. PCA was then applied to this matrix to determine all sixty-four principal components. Variance within object type and across object type were calculated for each iteration and compared. [21]. If the structure of the data was maintained across object types, we expect to see no difference between explained variance in the within or across task conditions.

## Results

### Single Feature Results

Single feature analyses were used to determine whether grasp or object accounted for a larger proportion of single-feature variance. Table 1 summarizes feature distribution across all three modulation categories. The majority of features were classified as modulating (either condition-independent or dependent) across both epochs. To control for multi-session concatenation, our statistical model included session day as a factor, as well as day interaction terms (day : grasp, day : object, day : grasp : object) to control interactions for inter-session variation. Representative day-invariant features modulating to grasp during the premovement and movement epochs are shown in **Figure 2a-2d** . No day-invariant features were found that modulated to object. The features shown in **Figure 2a-d** separated out by grasp and object are provided in ***Supplementary Figure 3*.**

**Figure 2:**
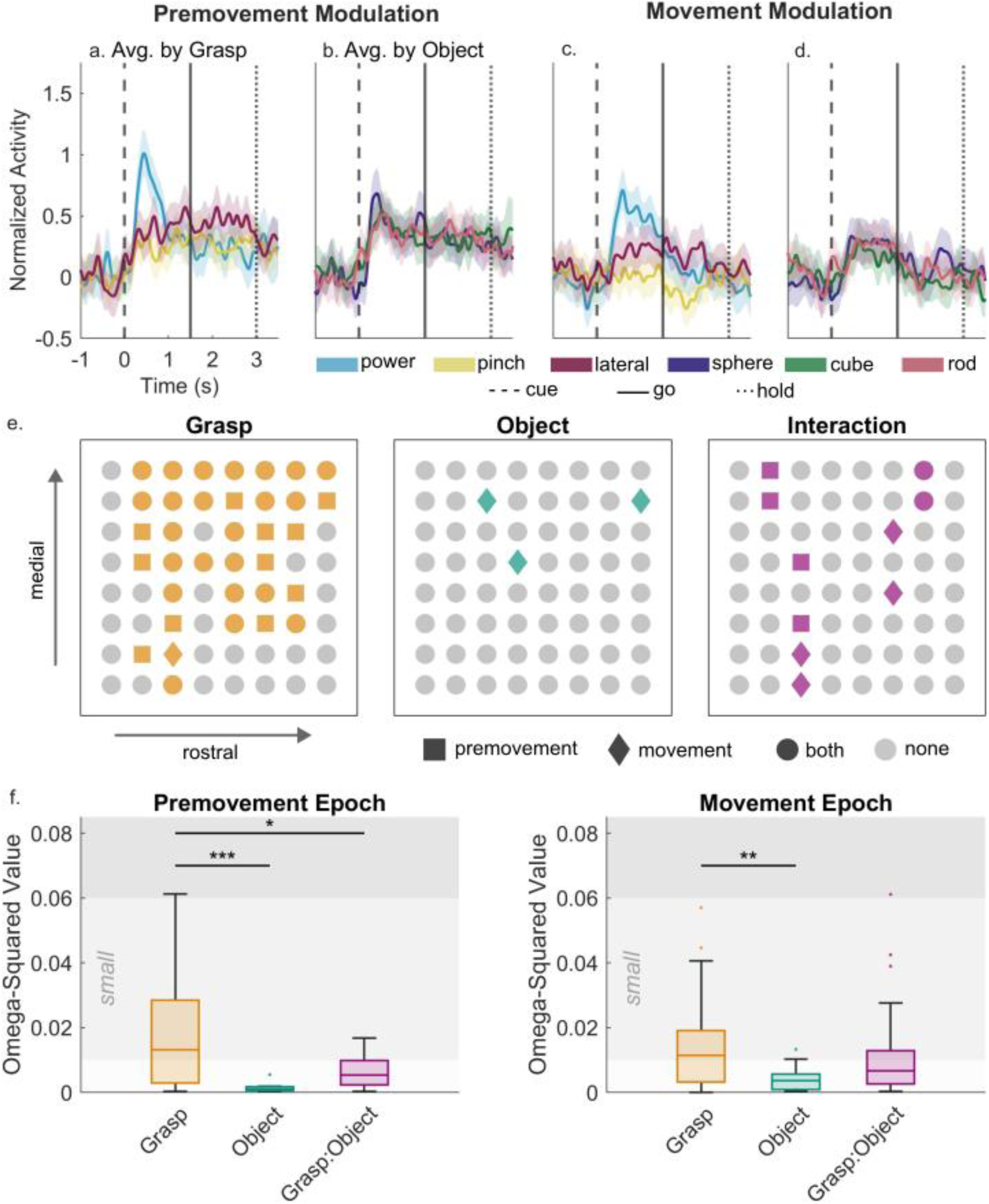
Sample Feature Traces and Modulation. Single feature traces (a-d) grouped by either grasp (left) shown with 95% confidence intervals that modulate to grasp in the premovement (a/b) and movement (c/d) epoch. For all single feature traces shown, vertical dashed line indicates cue (premovement) onset time, the solid line indicates movement onset, and the dotted line indicates the beginning of the hold epoch. e) Condition-dependent modulating features for grasp (orange), object (green), and pair type (purple), or none (grey) plotted based on their anatomical location on the array. For each modulating feature, those that modulate only in the premovement epoch have a square shape. Those that only modulate in the movement epoch have a diamond shape. Those that modulate in both epochs have a circle shape. Object modulating features were located slightly more rostrally than caudally on the array, while grasp and interaction modulating feature location was determined by which epoch it modulated in. f) Total omega-squared (ω²) metric for all features, regardless of modulation are plotted for grasp, object, and the interaction between grasp and object. Shaded grey regions indicate levels of significance for ω² values (ω² < 0.01: “negligible”, 0.01 < ω² < 0.06: “small”, 0.06 < ω² < 0.14: “medium”, ω² ≥ 0.14: “large”). For all comparisons, horizontal bars indicate significant difference between conditions as calculated from a 1-way ANOVA with Tukey-Kramer Corrections (p < 0.001, <0.01, and ≤ 0.05 indicated by ***, **, and * respectively).

**Table 1:** Modulation of Features (%) by Category. Features were categorized into non-modulating, condition independent, or condition dependent via a one-way and n-way ANOVA. Features were considered non-modulating if none of the nine conditions were found to be significantly different from baseline (p < 0.05). Condition-dependently modulating features were those that had at least one of the nine conditions significantly modulating from baseline, but no modulation to grasp, object, or the interaction between grasp and object. Features were only considered condition-dependently modulating if at least one condition modulated from baseline, and an n-way ANOVA with post-hoc corrections (Tukey’s HSD) found a significant difference between grasp pairs, object pairs, or grasp-object pairs. (p < 0.05)

|  | Non-Modulating | Condition Independent | Condition-dependent |
| --- | --- | --- | --- |
| Premovement | 6.3% | 53.1% | 40.6% |
| Movement | 10.9% | 40.6% | 48.4% |

We further evaluated features for tuning to grasp, object, or the interaction between grasp and object. The interaction term (grasp: object) evaluated whether the effect of grasp depends on the object, rather than testing for pairwise differences among the nine individual condition pairs. Across both epochs, condition dependent features predominantly modulated to grasp or the interaction between grasp and object. The multi-factor ANOVA revealed that in the premovement epoch, 53% of features modulated to grasp, 0% modulated to object, and 9% modulated to grasp-object interactions. These proportions changed in the movement epoch to 38% (grasp), 5% (object), and 9% (interaction). Crucially, all features that modulated to object or grasp-object interaction were found to also modulate to grasp. However, the reverse is not true as far more features responded to grasp alone than any other modulation category.

**Figure 2e** illustrates the spatial location of condition-dependent features on the array across the premovement epoch (square marker), movement epoch (diamond marker), or both epochs (circle marker). Non-modulating and condition-independent features are marked with a grey circle. Features that modulate to object were localized laterally and rostrally on the array. Premovement modulating features and features that modulated in both epochs tended to be more medial on the array, whereas grasp and interaction modulating features in the movement epoch were located more medially on the array.

Effect size, or the impact of each factor presented, was assessed via the omega-squared (ω²) metric across all 64 features, regardless of modulation category (**Figure 2f).** Each box represents the median, interquartile (IQR), and full range of values across all sixty-four features. Effect sizes showed that grasp had a significantly higher impact on variance in the data than object or interaction. During the premovement epoch, grasp values ranged from having a negligible to a medium impact (median 0.013, IQR 0.003-0.028) whereas object (median 0.001, IQR 0.0006-0.002) or grasp-object interaction (median 0.005, IQR 0.002-0.01) showed a negligible (0-0.01) to small impact (0.01-0.06). In the movement epoch, object (median 0.004, IQR 0.001-0.006) was found to have significantly smaller effect than grasp (median 0.011, IQR 0.003-0.02, 1-way ANOVA with post-hoc Tukey-Kramer corrections p < 0.05). No significant difference between grasp and grasp-object interaction (median 0.007, IQR 0.003-0.013) was observed.

Together, the single feature results demonstrate that grasp contributed the most to the variance in single feature data, followed by interaction. Object overall contributed a negligible amount to single feature variance. The impact of the interaction between grasp and object can also be seen on the single feature level.

### Population Results

To further assess the neural representation of grasp and object independently we computed the Euclidean distance between conditions within the reduced ten-dimensional latent space derived after aligning neural data across days (*See Methods)*. First, we assessed whether each condition was found to have significant above baseline modulation by calculating the average Euclidean distance between baseline activity and grasp-object pair activity **(Figure 3a).** Here, “pair” denotes the specific combination of grasp and object, not assessing whether the impact of grasp is dependent on object. Across both epochs, all nine grasp-object conditions were found to have above baseline Euclidean distances in 10-dimensional space. In addition, responses to each condition were variable across grasp and object pairs, demonstrating the impact of both grasp and object on the population level.

**Figure 3:**
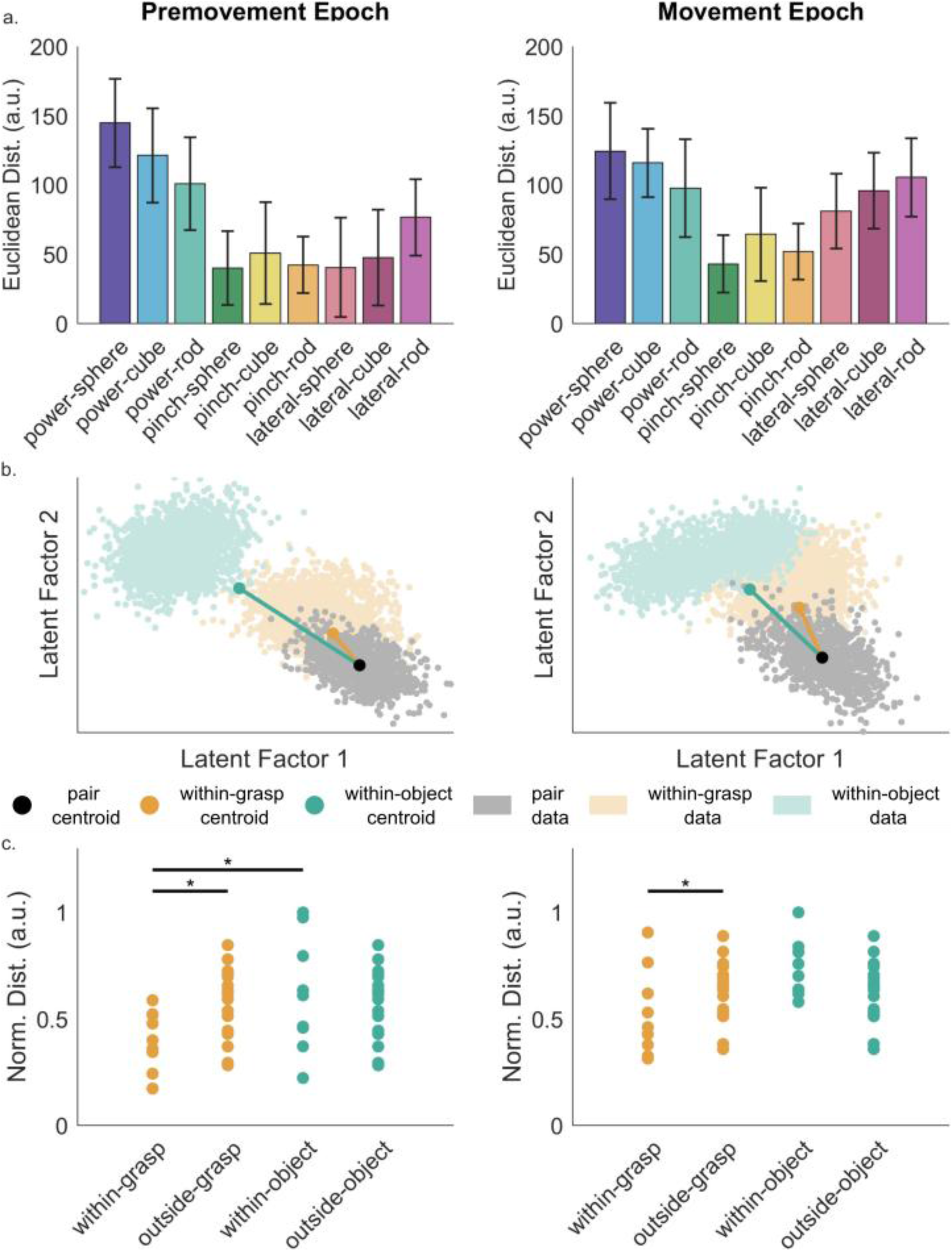
Euclidean Distance Between Conditions. a) Modulation depth as calculated by the Euclidean Distance between baseline activity and the grasp-object pair activity using a jackknife approach to generate 95% confidence intervals (grey bars). b) Sample distance metric visualization with bootstrapped data points projected onto the top two latent factors. For the sample pair (grey) the pairs of the same grasp (orange) and pairs of the same object (green) are plotted with the distance from the centroid (grey, orange, or green dot for the sample data, grasp data, and object data) of the sample data to the centroid of the grasp cluster and object cluster indicated with orange and green lines respectively. c) The normalized Euclidean distance between conditions grouped by within-grasp, outside-grasp, within-object, or outside-object with vertical bars indicating significance between groups (permutation test p < 0.05). Euclidean distance values were normalized such that maximum difference between conditions returned a value of 1 and minimum difference between conditions returned a value of 0.

To quantify whether the population structure is governed by grasp and object, we calculated the Euclidean distances between condition pairs. The normalized Euclidean distance between pairs of conditions is shown in **Supplementary Figure 4.** If population activity was driven strictly by grasp or object, we would expect values near zero within conditions sharing the same grasp or object respectively. Furthermore, if the population was overall more organized by grasp than object, pairs with the same grasp type should exhibit a smaller Euclidean distance than pairs with the same object (for example, pair type pinch-cube should be closer to pinch-rod than lateral-cube). To test this, each condition pair was categorized as a within-grasp (sharing the same grasp type), within-object (sharing the same object type), or outside-grasp and outside-object (different grasp and object types respectively) shown in **Figure 3c**.

Bootstrapped resampling (1,000 iterations) was performed for each of the nine pair conditions across both epochs and compared to a given sample pair. **Figure 3b**. illustrates this distance metric in a reduced two-dimensional latent space. Here, sample pair data (grey) is compared to pairs of the same grasp-type (within-grasp, orange) and pairs of the same object-type (within-object, green) with solid lines indicating the distances between condition centroids. This example shows on a reduced scale how the Euclidean distance metric is calculated and demonstrates how within-grasp distances vs. within-object distances are calculated. In this sample pair, the within-grasp distance was found to be less than the within-object distance. Distance comparisons for all nine conditions can be found in **Supplementary Figure 5 and Supplementary Figure 6.**

**Figure 3c** shows the resulting distances for all categories by taking only values above the diagonal of the 9x9 Euclidean distance matrix. Within-grasp distances were found to be significantly smaller than within-object distances in the premovement epoch only, but within-grasp distances were significantly smaller than outside-grasp values in both epochs. This indicates that condition pairs sharing the same grasp type occupy a tighter cluster in neural space than those sharing an object type. Thus, population level feature organization is governed primarily by grasp more than by object. While significance was not found comparing each of the nine pairs within-grasp values to within-object values, within-object values were consistently larger than within-grasp values on average for all nine conditions and both epochs (**Supplementary Figure 6**). However, pairwise Euclidean distances also provide clear evidence of population-level interaction effects. As previously stated, if the population strictly governed by grasp type, conditions sharing the same grasp type would project to nearly identical coordinates in state space, yielding within-grasp distances near zero. Instead, within-grasp distances exhibit consistently non-zero averages across both epochs (**Figure 3c**). This non-zero separation confirms that object also contributes to population activity organization, reflecting grasp-object interactions.

Assessing the population level separability of grasp (chance 33.33%), object (chance 33.33%), or pair (chance 11.11%), was achieved using 1,000 iterations of a 5-fold cross validated linear discriminant analysis model (**Figure 4**). Here, to evaluate grasp decoding, conditions with the same grasp type were pooled across object, and vice versa for object decoding. For comparison, decoding accuracies were calculated as percent above chance. Across both epochs, grasp classification yielded the highest decoding performance at 50.7±0.84% and 53.5±0.92% respectively. Grasp decoding accuracy was significantly higher than both object decoding (32.9±1.4% and 36.6±1.4%) in both epochs (**Figure 4b**). Pair above chance decoding was found to be comparable to grasp above chance decoding in the premovement epoch (28.5±0.85%), but significantly less than grasp in the movement epoch (28.2±0.97%). Comparing the confusion matrices for grasp and object decoding confirms that the population data is organized more strongly by grasp than object (**Figure 4C**). However, the 9-condition pair decoding matrix also provides a key insight into interaction effects. Similarly to the Euclidean distance metric presented earlier, if population activity was governed exclusively by grasp type, classification errors would be strictly confined within grasp types, collapsing over object dimensions. Instead, classification errors are not strictly bounded by grasp type. Instead, the diagonal values across individual conditions exhibit strong, above chance, classification in both epochs. This indicates that while grasp is the primary driver of population organization, interaction effects result in task-specific population codes for individual grasp-object combinations.

**Figure 4:**
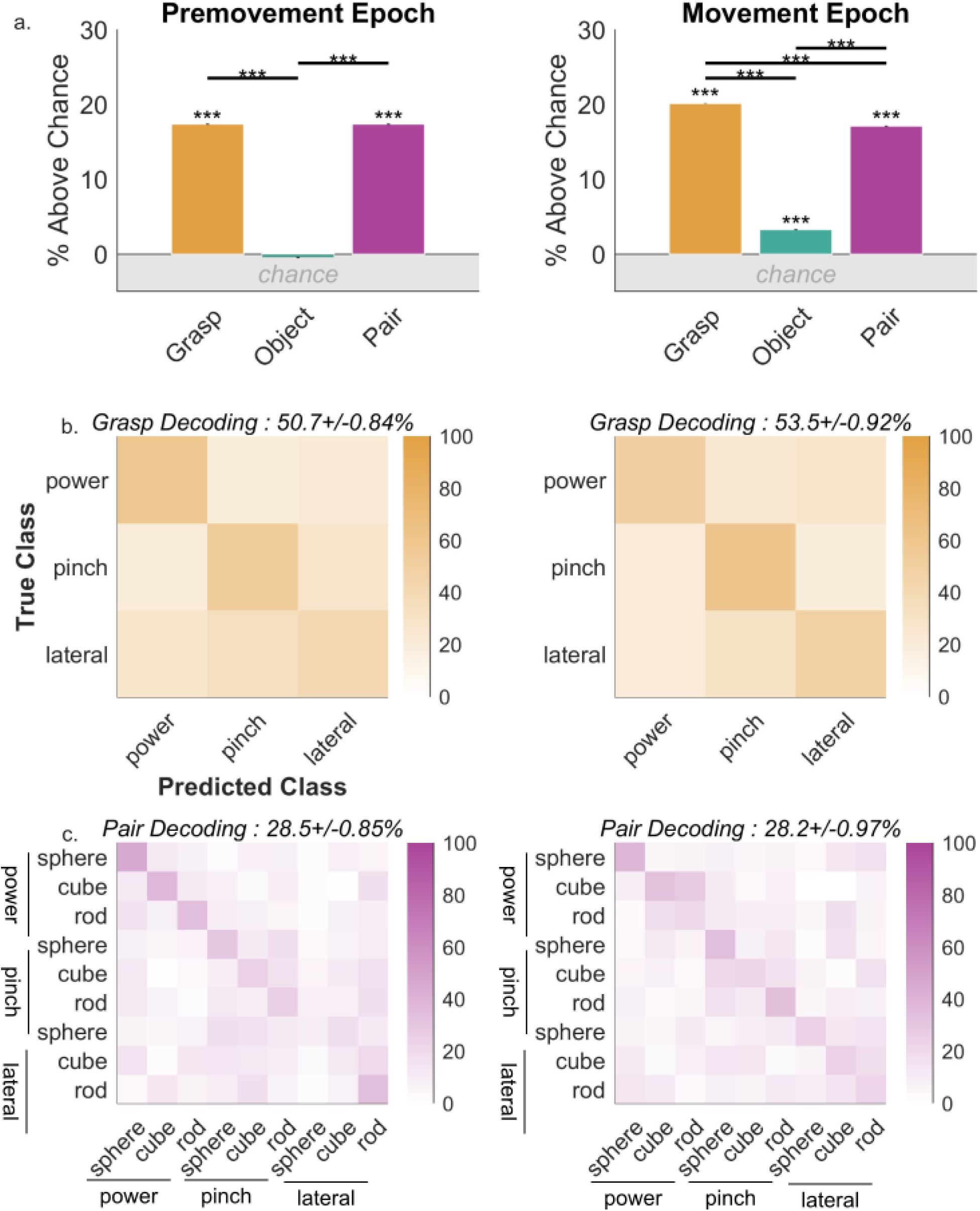
Decoding of Grasp, Object, and Pair. a) Decoding accuracies for grasp, object, and pair represented as percent above chance. Chance was calculated by shuffling the trial labels of the data set for 1000 iterations of 5-fold cross validated linear discriminant analysis. Vertical black bars indicate 95% confidence intervals. Asterisks above each bar indicate a significantly higher average decoding than chance (Welch t-test) or a significant difference between conditions with horizontal black bar (one-way ANOVA with Tukey-Kramer corrections). For both tests, p < 0.001 is indicated by *** respectively. Confusion matrices and cross validated decoding accuracy results of a 1000 iterations of a 5-fold cross validated linear discriminant model used to classify grasp (b) or pair (c).

While the decoding of grasp across both epochs were comparable, the patterns of decoding were slightly different. During the premovement epoch, power and pinch were mostly highly distinguishable from each another, whereas during movement, all three grasp types were found to be distinguishable from one another above chance (**Figure 4a**). Pair decoding successfully classified the majority of the pairs above chance for both epochs. However, in the movement epoch, off-diagonal conditions within the same grasp category increased, reflecting a decrease in object separability within a given grasp type. For instance, in the premovement epoch, power conditions were decodable from one another at 46±1.8%,38±0.3%, 34±3.6% for sphere, cube, and rod. In the movement epoch, these diagonal values decreased to 39±2.1%, 33±3.3, and 21±0.3%, with the off diagonal misclassifications reaching up to 28±2.9%. Similar trends were observed in pinch and lateral grasps. This epoch-dependent shift indicates that while planning maintains a distinct object separation within a grasp, movement execution drives the population towards grasp-level clustering.

The same boot-strap iteration method mentioned above for generating sample pair data for **Figure 3B** was used to generate samples of the time-varying pair trajectories for all ten latent factors. The resulting data was averaged and plotted with 95% confidence intervals shown in **Figure 5a-5d**. The trial-averaged neural trajectory across all conditions and iterations was first calculated to identify the condition independent signal (**Figure 5a**). The condition independent trajectory was then subtracted from each pair iteration to amplify the variance pair condition variance (**Figure 5b**). Trajectories were then either averaged over grasp (**Figure 5c**) or object (**Figure 5d**) to visualize condition spread within the first latent factor. The variance between grasp, object, and pair conditions were calculated (**Figure 5e**) and compared to one another. Within the first latent factor, the variance due to grasp was substantially higher in magnitude than the variance due to object. Furthermore, the 9-class pair variance trajectory closely mirrored the grasp variance trajectory throughout the trial duration. This indicates that within the primary latent factor, accounting for the largest proportion of population variance, grasp serves as the dominant driver of condition separation. Secondary latent dimensions (***Supplementary Figure 7***), accounting for overall less variance, revealed a greater separation between object types. This confirms that while interaction effects shape population trajectories, they operate a secondary organizational axis relative to grasp. Analyzing the temporal profiles of the variance traces revealed key differences in feature processing across epochs. The times of peak variance within the premovement epoch were found to be similar for grasp and pair variance at 492.6 ± 26.5ms and 485.7 ± 48.2ms after cue time. However, peak object variance was found to be at 181.15 ± 157.8ms. This temporal offset indicates that while object exerts a weaker overall effect on population variance, object feature information is processed earlier during premovement planning before integration into the grasp dominated population.

**Figure 5:**
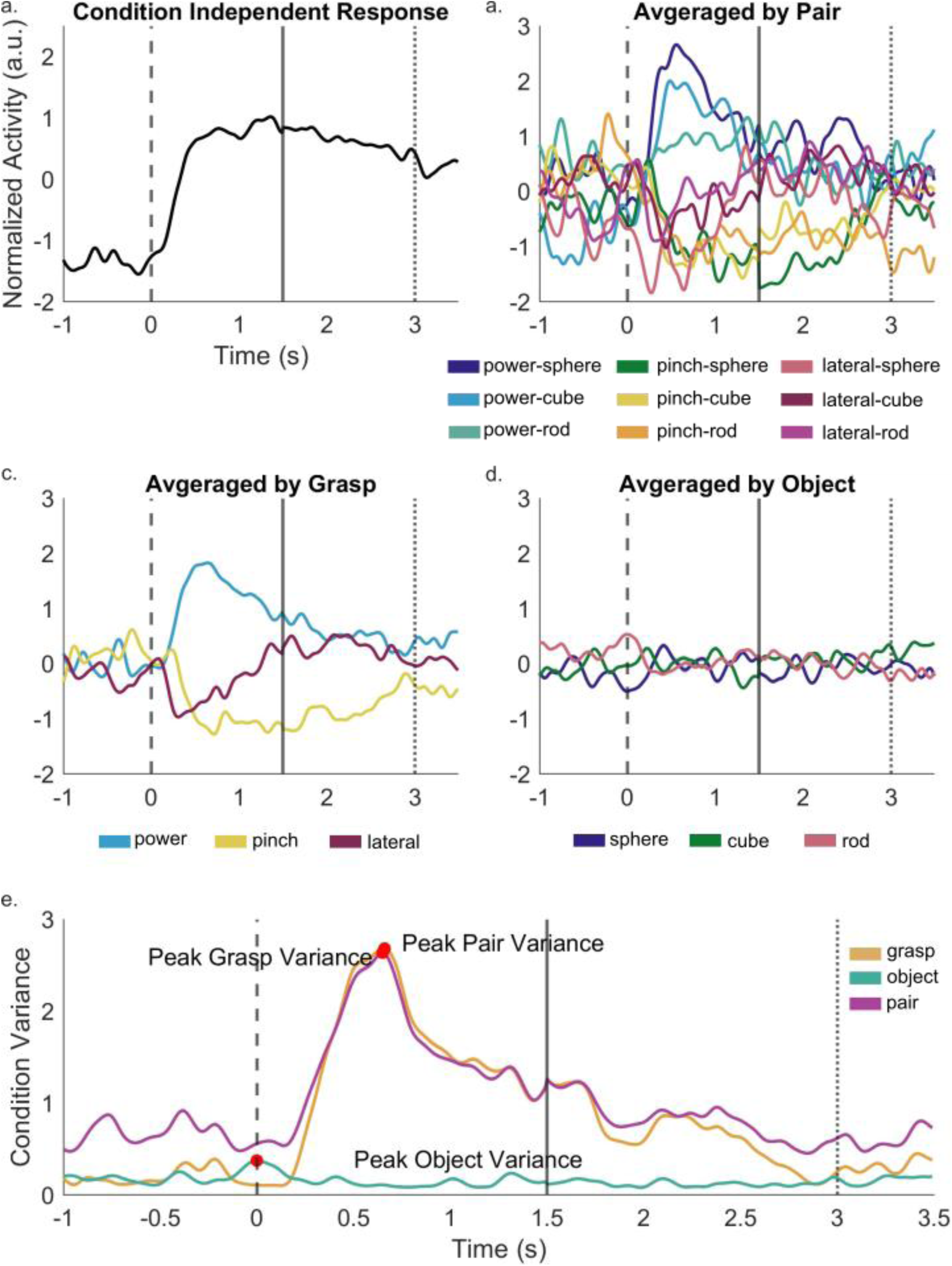
Time Varying Projections of Pair Conditions on First Principal Component. Random samples of the data were selected for 1000 iterations and then averaged over the nine individual conditions. a) Time-varying projections of the nine conditions were averaged to reveal the time-varying signal underlying all pair data. b) The time-varying projection was then subtracted from each of the nine pair conditions to highlight the differences between pair type with 95% confidence intervals. Data was then averaged over grasp type (c) or object type (d) and plotted with 95% confidence intervals. e) The variance between grasp, object, and pair types. Higher variance over time indicates higher separability of the conditions. Peak variance within the premovement epoch is noted with a red dot and corresponding labels. For all traces shown, dashed line indicates cue (premovement) onset time, the solid line indicates movement onset, and the dotted line indicates the beginning of the hold epoch

While grasp is the main factor contributing to data variance in IFG, previous results (**Figure 2f**) indicated the effect of grasp-object interactions on single feature and population level variance. Within-grasp object decoding revealed that object discriminability is strongly dependent on the corresponding grasp type

(**Figure 6a**). Across both epochs, power grasp yielded the highest overall object decoding accuracy followed by pinch and lateral grips (p<0.05). These results closely mirrors the participant’s subjective selection of optimal grasp-object pairings (**Table 2**). When presented with individual object, the participant ranked power grips the most natural choice for spheres and cubes, but selected the lateral grip as the most natural choice for rods.

**Figure 6:**
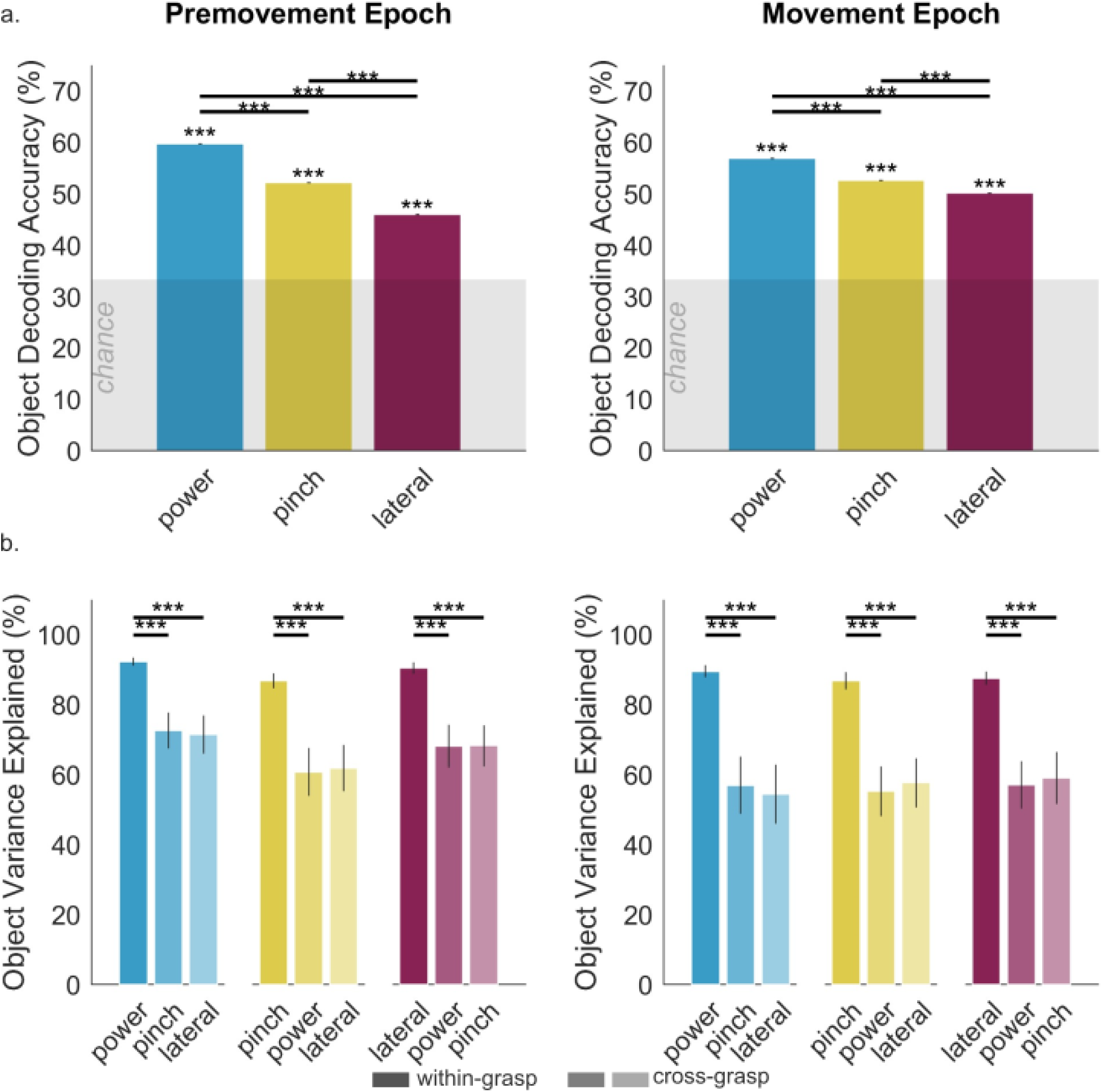
Impact of Object Type on Neural Data Structure. Cross validated 5-fold linear discriminant analysis was performed for 1,000 iterations to decode grasp across each of the object types presented. The decoding accuracy for the premovement epoch and movement epoch shown are shown with asterixis above the bars to indicate that decoding accuracy was significantly above chance. Vertical bars indicate 95% confidence intervals. Shaded grey area indicates chance. b) Premovement and movement epoch neural data of a held grasp condition was projected onto the first five principal components subspace. Darker bars (within-object) indicate variance accounted for when projecting the object neural data onto its own top five principal components and lighter bars (cross-object) indicate projecting onto a different object’s top five principal components. Vertical bars indicate standard deviation. Horizontal bars indicate significant differences between conditions as determined by a one-way ANOVA with post-hoc Tukey-Kramer corrections (a and c) or for a two-sided t-test (b and d). p < 0.001, <0.01, and ≤ 0.05 indicated by ***, **, and * respectively.

**Table 2:** Participant Selectivity Responses. RP1 was presented with all three object individually as shown in the visualization and asked to rank the three grasp types from most likely to least likely to use for the given object. The survey was also done where the grasp type was presented and asked to rank objects from most likely to least likely to use with the given grasp. The participant was not given other instructions other than the ones provided above

|  | *Most Likely → Least Likely* |  |  |
| --- | --- | --- | --- |
| *Object Condition* |  |  |  |
| sphere | power | pinch | lateral |
| cube | power | pinch | lateral |
| rod | lateral | power | pinch |
| *Grasp Condition* |  |  |  |
| power | sphere | cube | rod |
| pinch | cube | rod | sphere |
| lateral | rod | sphere | cube |

Inspection of the within-grasp object decoding confusion matrices confirmed that neural state-pace organization during the premovement reflects the participant’s subjective optimal grasp-object pairings (**Supplementary Figure 8**). When presented with individual grasps, the participants’ preferred pairings were power-sphere, pinch-cube, and lateral-rod. In the pre-movement epoch, the highest-performing object decoding within each grasp type corresponded exactly to the participant rankings with power-sphere (63±2%), pinch-cube (56±2.1%), and lateral-rod (59±3.1%). Conversely, movement execution resulted in a shift of object-separability patterns. While power-sphere remained a high performer (71±2%), decoding accuracies for pinch and lateral grasp shifted to pinch-sphere (61±3.5%), and lateral-cube (55±2.9%) during the movement epoch. Interestingly, these combinations were rated as the least likely pairings in the subjective survey.

Quantification of the manifold alignment across conditions was assessed through the object activity variance accounted for (VAF) within and across grasp types (**Figure 6b**). Neural trajectories for a given grasp were projected onto the top five principal components derived from either the same grasp (within-grasp) or a different grasp (cross-grasp). Across both the premovement and movement epochs, cross-grasp VAF (**Figure 6b**, second and third bar per grasp group) was significantly lower than the within-grasp VAF (first column of every grasp-type, p < 0.05). These results demonstrate a significant difference in the underlying neural-manifolds of each grasp type. Together, these results confirm that while grasp type drives population level organization, object type alters the within-grasp state space.

## Discussion

The primary finding of this study is that grasp serves as the principal driver of neural variance in human inferior frontal gyrus (IFG), far outweighing the individual contribution of object. While previous microelectrode array recordings in a person with tetraplegia demonstrated that specific grasp-object pairings can be decoded from premotor areas, specifically ventral premotor cortex, during motor imagery [12], our findings provides additional mechanistic insight into whether the object type or the grasp type is the primary driver of modulation. Our population level results align directly with human EEG and fMRI studies that attributed the function of human IFG to be encoding grasp parameters, and not object parameters [22], [23]. At the single feature level, while the predominant number of features recorded modulated to grasp type, a subset of features modulated to object and the interaction between grasp and object. Crucially, we found that all features that were selective for object type and grasp-object interaction were also selective for grasp type. This hierarchy reflects mixed selectivity within IFG, mirroring single unit studies in macaque area F5 [4], [24].

Although grasp configuration is the primary driver of population organization, object remains distinctly separable within individual grasps through interaction terms. First, we found that the temporal trajectory analyses revealed that object variance peaks approximately 200ms earlier in the premovement epoch than grasp or pair variance (∼180ms vs. ∼490ms post-cue). This mirrors non-human primate studies where F5 ensembles process visual object features early in the delay period before transforming them into a specific motor plan [3], [24], [25]. In addition, canonical neurons activating to specific grasp-object pairings dictate how objects are separated within a given grasp. These studies demonstrate that early visual input selectivity activates canonical neuron populations based on habitual grasp-object pairings that configure the state space prior to movement execution [2], [4], [26]. This separability is demonstrated in **Figure 6a**. Importantly, because object position and rotation were fixed within each grasp condition, held-condition object decoding confirms that this within-grasp separability reflects intrinsic sensorimotor mapping rather than spatial location or orientation of the object. Together, these results agree with previous literature that while grasp is the principal driver of IFG modulation, interaction terms heavily shape the specific patterns of object separability within a grasp type.

Assessing the functional role of the separation between objects within a grasp relies heavily on the mechanisms of premotor canonical neurons. Previous studies have established that canonical neurons exhibit selective tuning for congruent grasp-object pairs, firing when an observed object matches the preferred grasp, regardless of if a movement occurs [4]. This automatic visual-motor mapping explains why in the premovement epoch, the selection of optimal objects within in a specific grasp type aligns exactly with the participants survey responses. In addition, subspace alignment analysis provided further proof of grasp-dependent state space reorganization (**Figure 6b**). When neural data of a given grasp were projected onto the top five principal components derived from another object type, the variance accounted for was found to be significantly lower than when projecting the neural data onto the within-grasp manifold (p<0.05) for all combinations. The differences in manifold structure indicates the impact that grasp-object interactions have on the neural data, demonstrating that each grasp has its own individual mechanism for separating object types. However, in cross-grasp conditions there is a substantial, non-zero variance accounted for, indicating a partially shared underlying neural manifold structure. This is to be expected due to the fact that we compared two manifolds generated by grasping tasks and patterns of activity used across grasps tasks should have some degree of similarity. This balance of shared grasp configuration but significantly different object organization further confirms that the primary driver of population level activity is by grasp type, but still affected secondarily by grasp-object interactions.

An intriguing finding from this study is the dynamic shift in state-space organization between the premovement and movement epoch. While IFG is primarily characterized as a premovement area, we initially used the movement epoch as a comparison to the premovement epoch to verify early premovement tuning. However, population decoding remained high within both epochs but with distinctly different state-space structures. For example, in the premovement epoch, power and pinch grasps were most separable in grasp decoding, whereas all three grasp became comparably separable during the movement epoch. This temporal reorganization is consistent with sub-populations of neurons identified in non-human primate literature which characterize area F5 neurons as sensory, sensory motor, or motor based on their response profiles. Non-human primate literature subdivides area F5 into F5c, F5p and F5a [27]. F5a is more connected to sensory and parietal areas for the integration of object information [28], [29], whereas F5c and F5p are activated during hand and mouth movements respectively [30]. Spatial mapping of our microelectrode revealed a similar anatomical gradient. Premovement modulating features were clustered more laterally on the array, whereas features modulating exclusively during the movement epoch clustered more medially on the array. This spatial distribution suggests that lateral premovement features reflect early visual-motor integration, responding to both object properties and grasp type properties [4], [5], [6], [7], [8] whereas medially clustered movement modulating features are predominantly modulated by grasp type alone [3].

This movement epoch recruitment of dedicated motor features coincides with a reduced impact of grasp-object interactions. In the pair decoding matrix (**Figure 4b**), within-object misclassifications increased during the movement epoch, indicating that object separation within a grasp type is reduced. This finding is comparable to the findings of Fluet. Et. al.[25], where the population activity of area F5 reflected the grasp type used more so than the object properties during the movement execution phase. The interaction suppression reconciles why the least likely pairings in our behavioral survey exhibited the highest decoding discriminability during movement (**Supplementary Figure 8**). Because IFG canonical and motor ensembles display preference, or an increased firing rate, for specific visual motor pairings, presenting a non-ideal or biomechanically incongruent pairing can result in a firing-rate suppression or reduced population activity relative to optimal pairings [25]. As confirmed in **Supplementary Figure 9,** non-ideal pairings of pinch-sphere (column 2, green trace) and lateral-cube (column 3, light pink trace) have reduced activity in comparison to the other pairs within the same grasp in the movement epoch. The suppression of activity in turn results in a larger separation between grasp-modulating canonical neurons and the non-ideal pairing, allowing linear classifiers to achieve high separability of the non-ideal pairing, despite being driven by population activity suppression.

Overall, this study demonstrates that grasp type is highly represented in IFG, regardless of object type presented. IFG presents an advantageous target for BMI systems due to the high separability between grasp types before movement onset occurs [12]. In contrast, traditional decoders built on M1 information have focused on decoding hand kinematics during movement [31], [32], [33], [34], [35], [36]. In addition, we found that object types are separable within a grasp likely driven by canonical neuron activity.

However, while interaction effects alters the grasp state space, or the separation between the object types, it does not collapse population level grasp separability. These organization principles offer key insights into the design of future BMI decoders utilizing IFG neural information.

## Acknowledgements

The authors wish to thank the study participant and their family for their time and commitment to this study.

## Author Contributions Statement

E. C.C and A.B.A. designed the study. E.C.C. performed the experiments, analyzed the data, wrote the main manuscript, and compiled the Supplementary Information. C.F. assisted in data analysis and writing for the cross-projection analysis. W.M. assisted in data collection. A.B.A supervised data collection and guided the project. J.S. and E.H. planned and performed the microelectrode array surgery. All authors reviewed and edited the manuscript.

## Data Availability Statement

The data can be made available upon reasonable request by contacting the lead or senior authors.

