## Supplementary Figures for "Assessing the Relative Impact of Grasp and Object on Inferior Frontal Gyrus Activity during a Grasping Task"

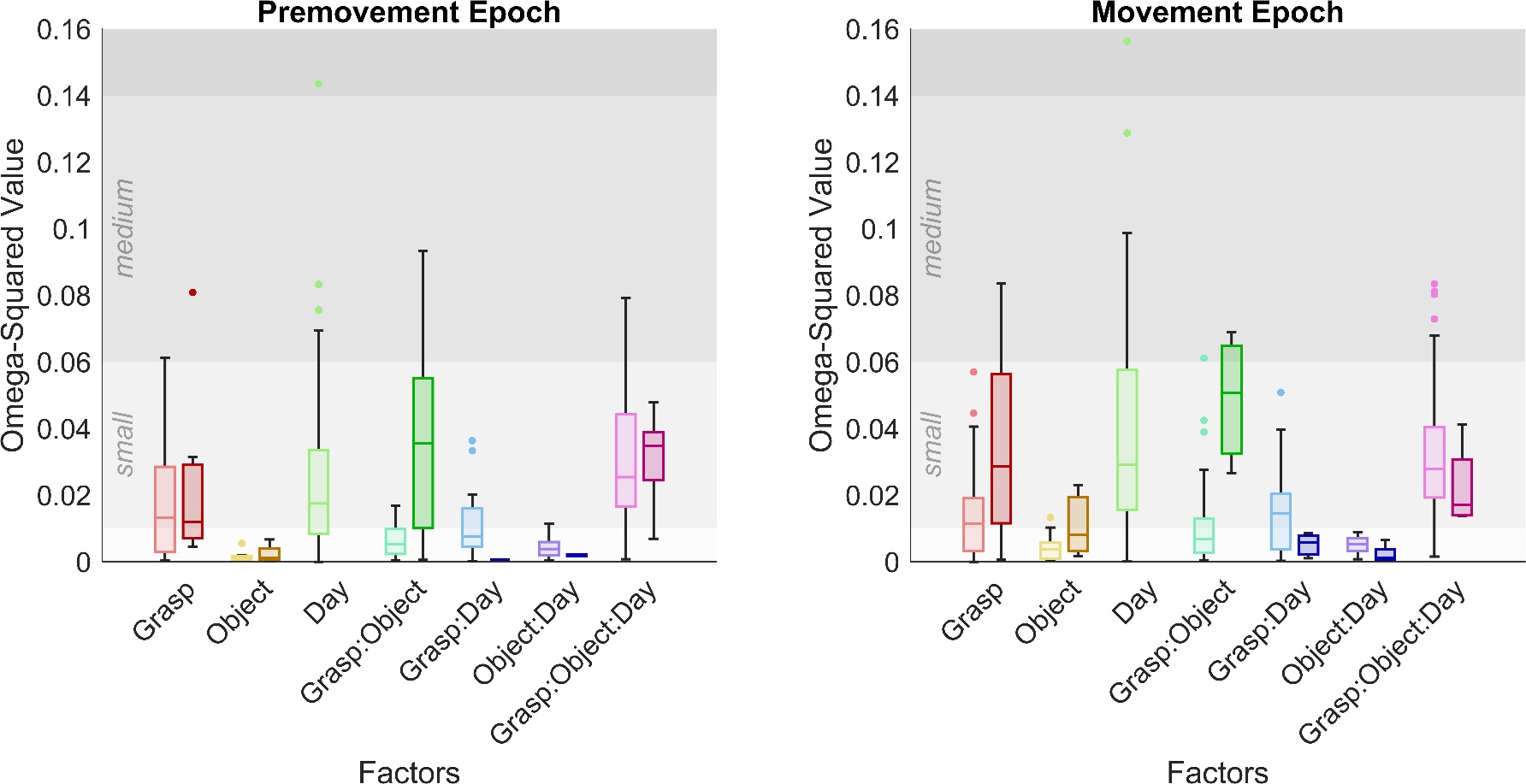

***Supplementary Figure 1: Impact of Factors on Single Feature Data Calculated by Omega-Squared Reduction of Impact of Day-Related Factors on Omega-Squared (ω²)*** ***Values Post-Alignment.*** *Total omega-squared (ω²) metric for all features, regardless of modulation are plotted for grasp, object, and the interaction between grasp and object. Shaded grey regions indicate levels of significance for ω² values (ω² < 0.01: “negligible”, 0.01 < ω² < 0.06: “small”, 0.06 < ω² < 0.14: “medium”, ω² ≥ 0.14: “large”). Lighter colors indicate ω² values before alignment and darker colors indicate ω² values after alignment. Almost all factors regarding day (Day, Grasp: Day, Object: Day) had lower ω² values after alignment. Grasp : object : day interactions were found to be increased in median value for the premovement epoch but reduced in the movement epoch and an overall reduction in the range of values for both epochs. Post-alignment values all fell into the negligible to small range. In addition, grasp, object, and grasp : object interactions were found to increase post-alignment.*

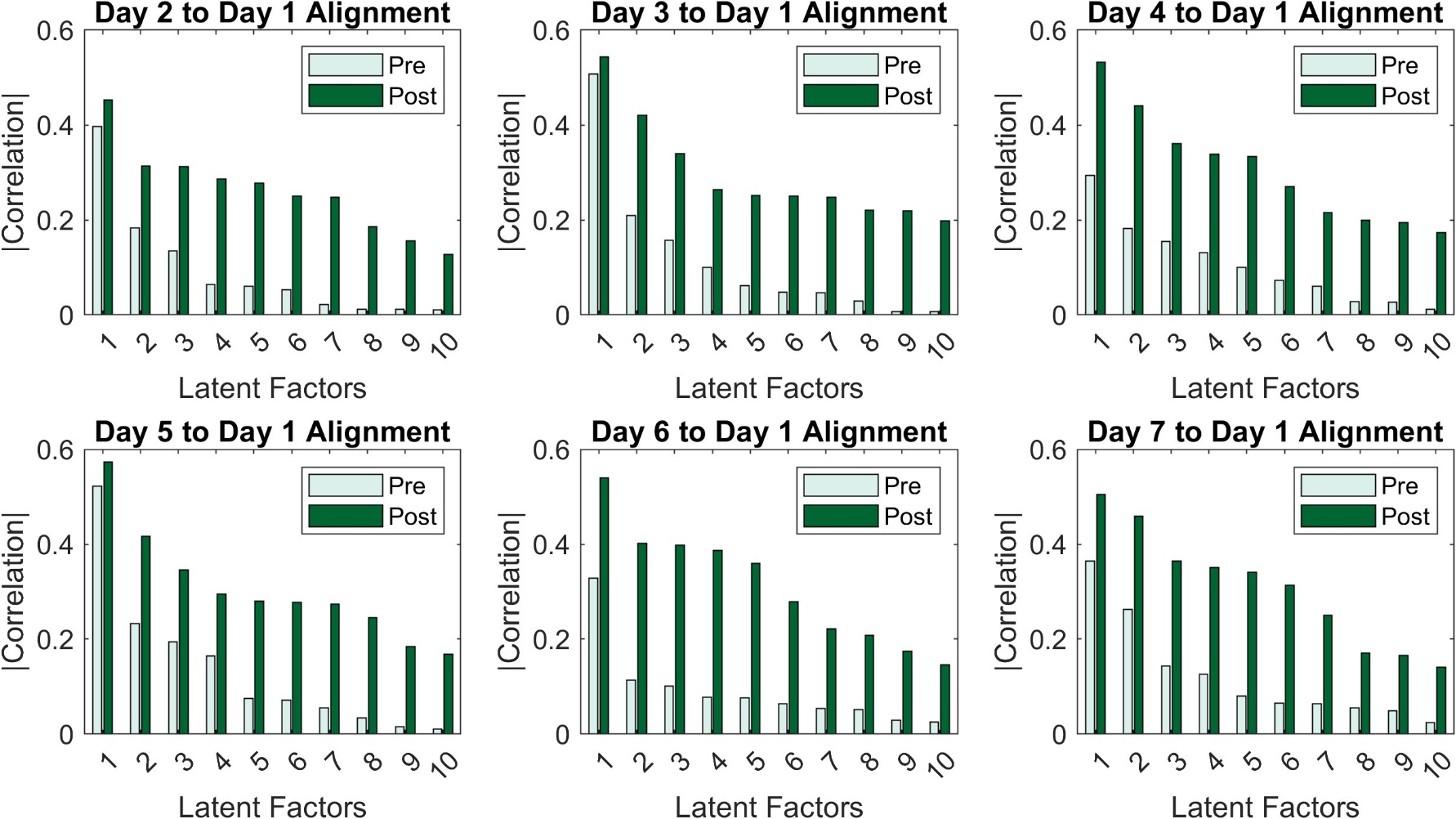

***Supplementary Figure 2: Alignment of Latent Factors by Day.*** *Principal component was applied to identify the coefficients and score of the top ten latent factors in condition averaged data. This coefficient was then used to project individual trials into reduced dimensional space. Procrustes alignment without scaling or reflection was applied to identify the transformation from reference latent factors (Day1) to all other days of data. This transformation was then applied to the individual trials resulting in a matrix maintaining the original trial count but with a reduced number of “features” representing the top ten latent factors. Correlation between the condition averaged latent factors was calculated and plotted for each day and factor. Light green represents the correlation between the days before alignment and dark green represents the correlation between days after alignment.*

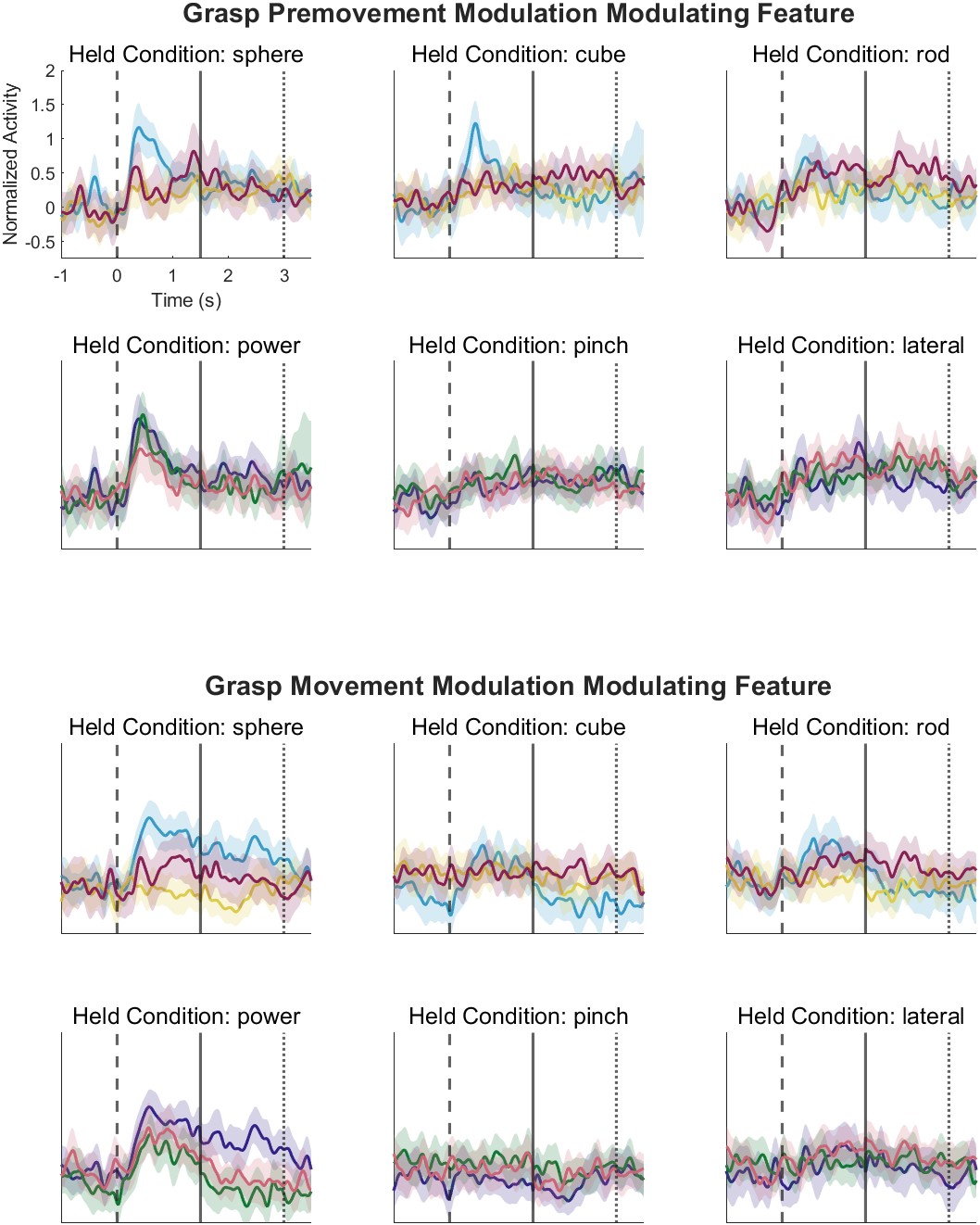
***Supplementary Figure 3: Sample Single Feature Modulation Separated out by Object and Grasp Type.*** *The same features shown in Figure 2a and 2b, separated by held condition. Traces are plotted with 95% confidence intervals*

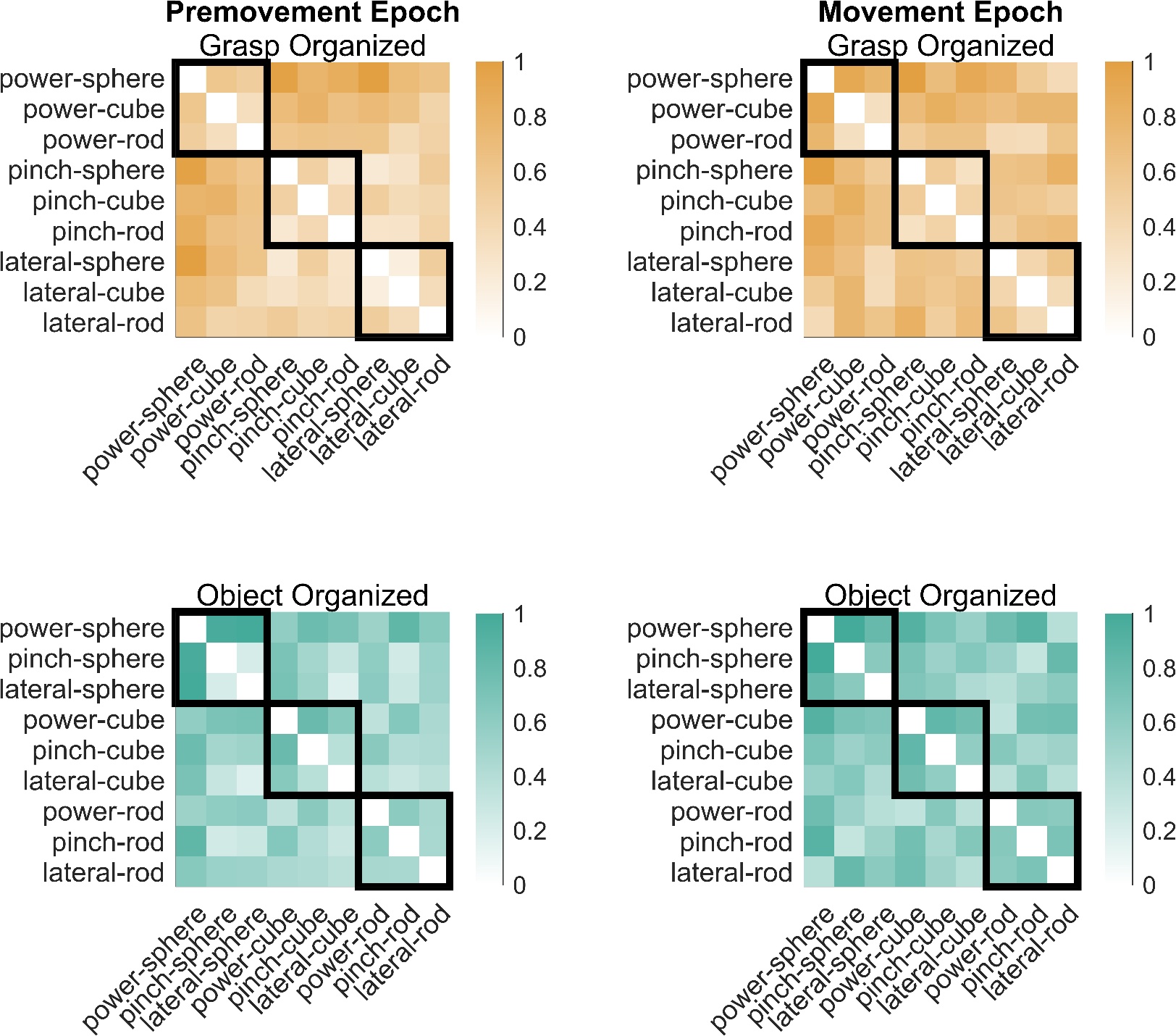

***Supplementary Figure 4: Full Normalized Euclidian Distance Confusion Matrix.*** *Cross-validated Euclidean Distance was calculated between each pair of data and normalized by the maximum distance found between pairs. Grasp organized and object organized matricies contain the same results but ordered either by grasp type or object type.*

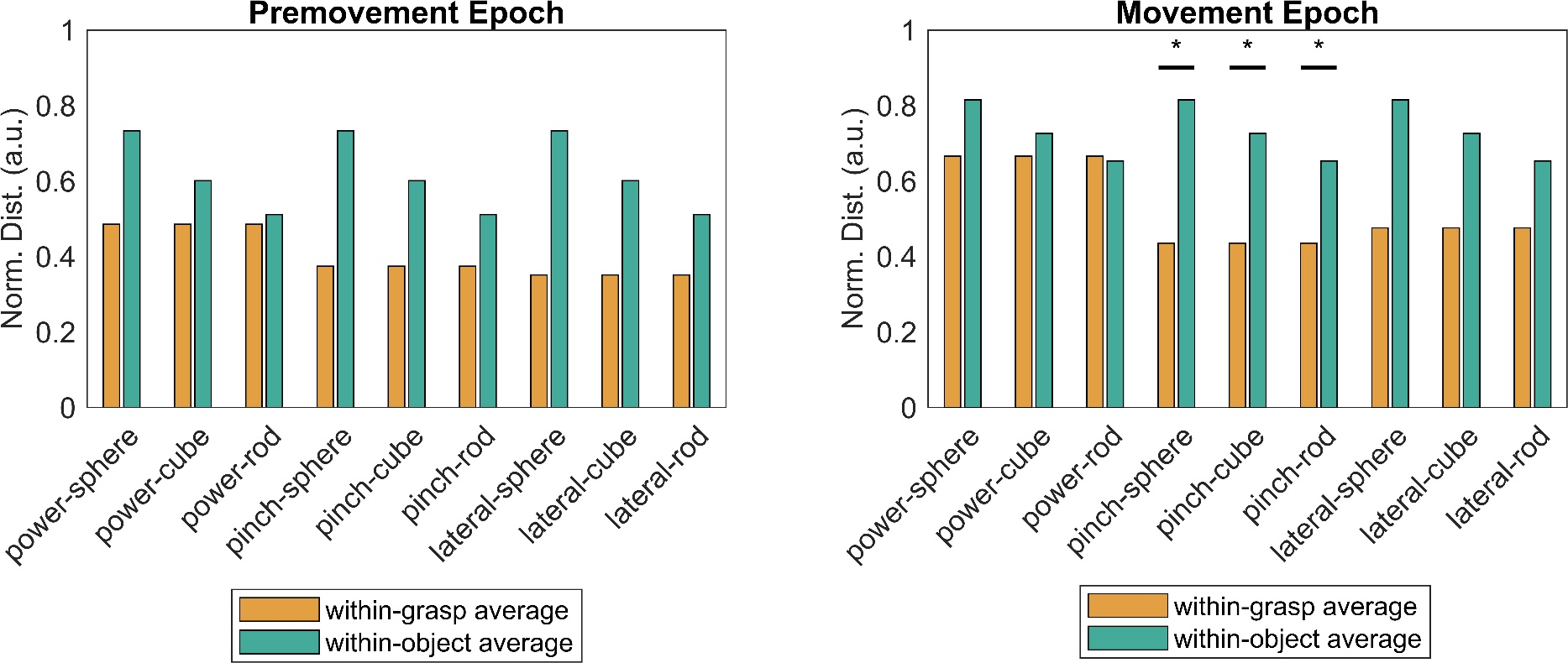

***Supplementary Figure 5: Comparison of Within-Grasp to Within-Object Values by Pair****. For each pair, the normalized Euclidan Distance for pairs with the same grasp type (within-grasp) and pairs with the same object type (within-object) were compared with a two sided t-test (p<0.05). Three values for each category were compared with only few pairs resulting in significance.*

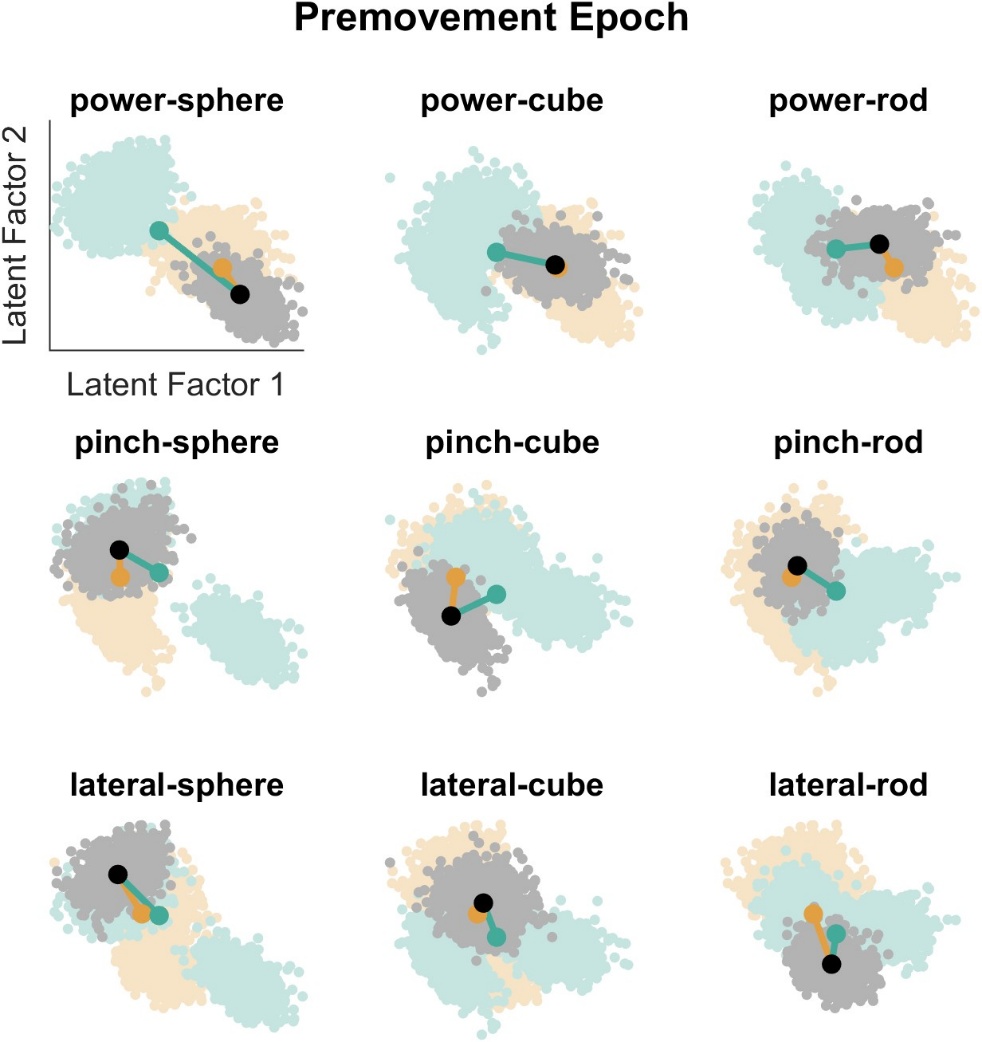

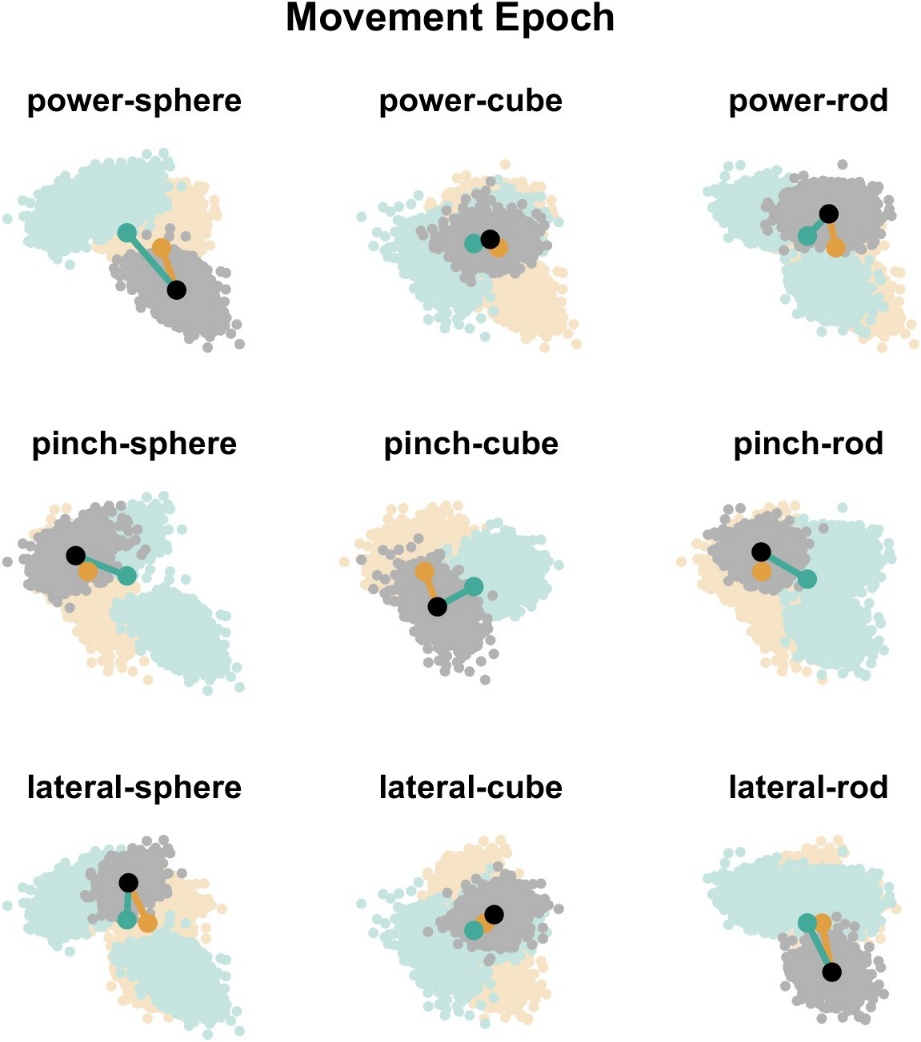

***Supplementary Figure 6: Sample Bootstrapped Data Plotted onto Latent Factor 1 and Latent Factor 2.*** *A 1,000 iteration bootstrapped resampling procedure was done to generate a sample data set where in each iteration, a random selection of the data within each condition was sampled and averaged. Each point representes a single iteration. Sample distance metric visualization with bootstrapped data points projected onto the top two latent factors. For the sample pair (grey) the pairs of the same grasp (orange) and pairs of the same object (green) are plotted with the distance from the centroid (grey, orange, or green dot for the sample data, grasp data, and object data) of the sample data to the centroid of the grasp cluster and object cluster indicated with orange and green lines respectively. It is important to note that the centroids include the position of the grey cluster when calculating the distance line, as it belongs to that grasp or object type.*

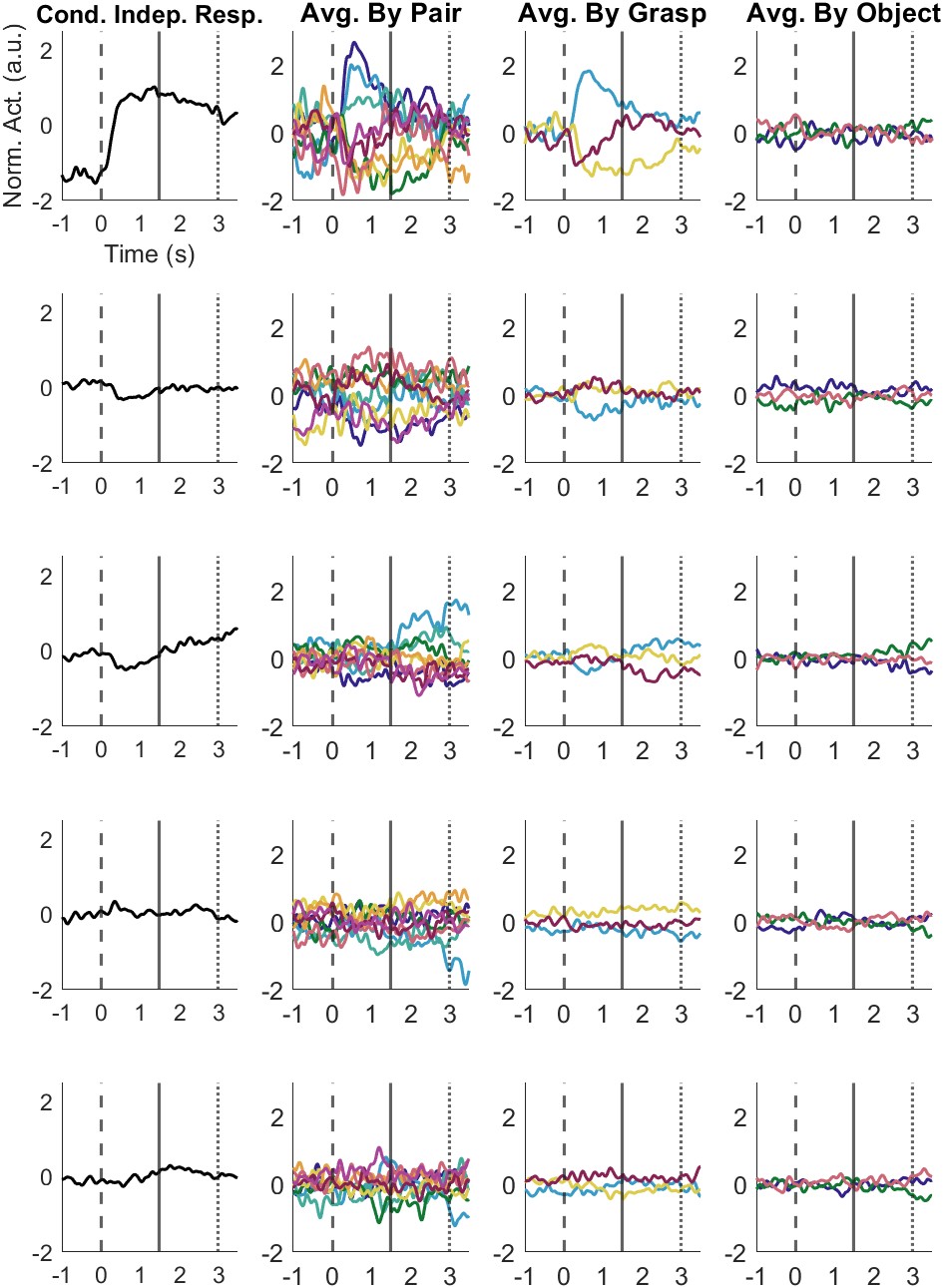
***Supplementary Figure 7: Additional Latent Factor Projections.*** *Latent Factor 1-5 time-varying projections to identify the condition independent response, averaged by pair response, averaged by grasp response, and averaged by object response as described in Figure 5.*

*Latent Factor 5*

*Latent Factor 4*

*Latent Factor 1*

*Latent Factor 2*

*Latent Factor 3*

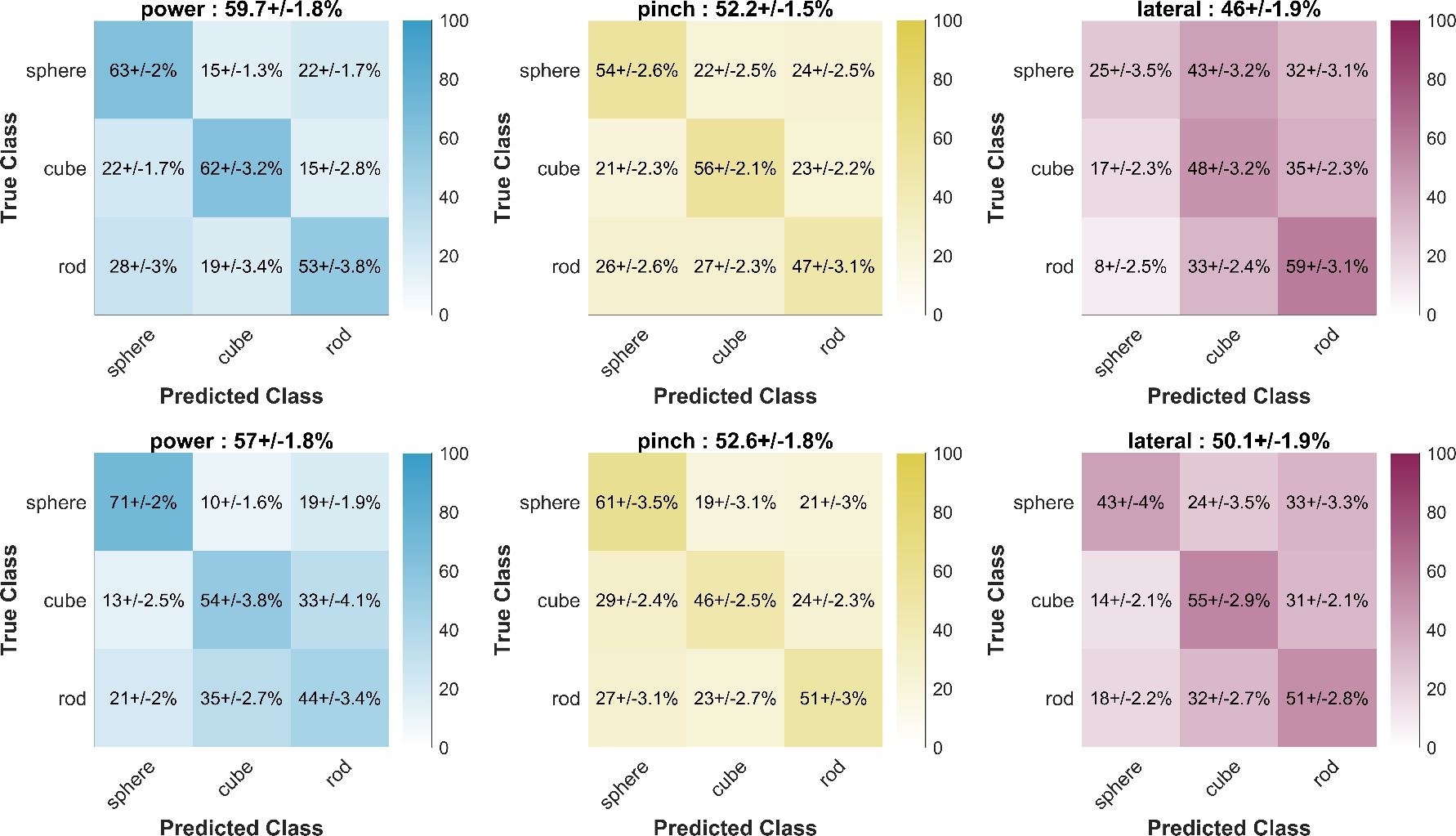

**Movement Epoch**

**Premovement Epoch**

***Supplementary Figure 8: Confusion Matrices by Held Grasp Condition.*** *As outlined in the methods of Figure 4 and Figure 6, a 1,000 iterations of a 5-fold cross validated Linear Discriminant Analysis model was used to identify the separation between object conditions within a specific grasp. Values overlaying each cell of the confusion matrix is mean +/- standard deviation across the 1,000 iterations.*

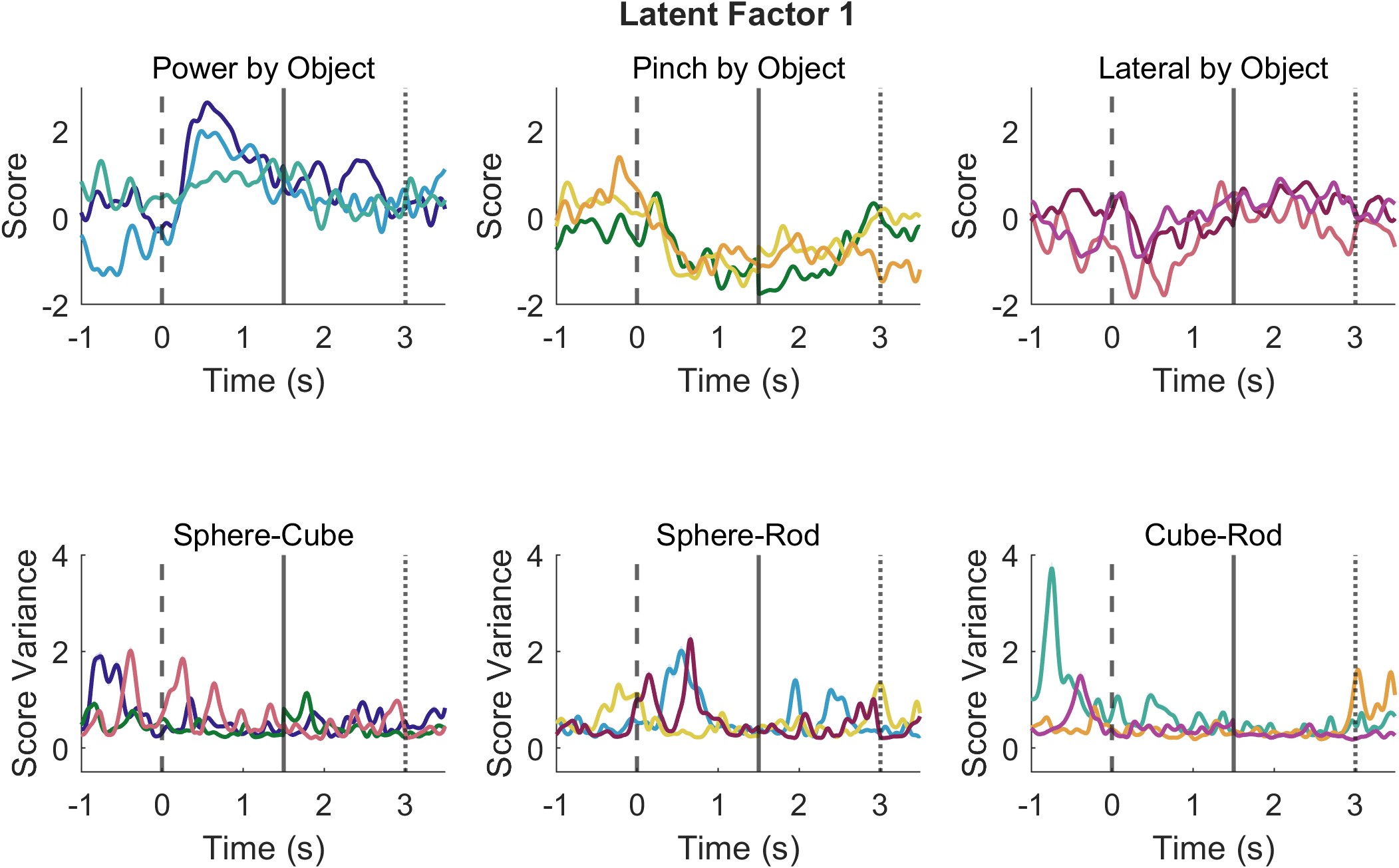

***Supplementary Figure 9: Separation of Latent Factor Projections by Grasp Type****. The same methodology used in Figure 6 and Supplementary Figure 7 was used to separate each of the nine pair conditions by their object type.*
